# Quantifying peripheral modulation of olfaction by trigeminal agonists

**DOI:** 10.1101/2023.03.13.532477

**Authors:** Genovese Federica, Xu Jiang, Tizzano Marco, Reisert Johannes

**Affiliations:** Monell Chemical Senses Center, 19104 Philadelphia, PA, USA

## Abstract

In the mammalian nose, two chemosensory systems, the trigeminal and the olfactory mediate the detection of volatile chemicals. Most odorants in fact are able to activate the trigeminal system, and vice versa, most trigeminal agonists activate the olfactory system as well. Although these two systems constitute two separate sensory modalities, trigeminal activation modulates the neural representation of an odor. The mechanisms behind the modulation of olfactory response by trigeminal activation are still poorly understood. In this study, we addressed this question by looking at the olfactory epithelium, where olfactory sensory neurons and trigeminal sensory fibers co-localize and where the olfactory signal is generated. We characterize the trigeminal activation in response to five different odorants by measuring intracellular Ca^2+^ changes from primary cultures of trigeminal neurons (TGNs). We also measured responses from mice lacking TRPA1 and TRPV1 channels known to mediate some trigeminal responses. Next, we tested how trigeminal activation affects the olfactory response in the olfactory epithelium using electro-olfactogram (EOG) recordings from WT and TRPA1/V1-KO mice. The trigeminal modulation of the olfactory response was determined by measuring responses to the odorant, 2-phenylethanol (PEA), an odorant with little trigeminal potency after stimulation with a trigeminal agonist. Trigeminal agonists induced a decrease in the EOG response to PEA, which depended on the level of TRPA1 and TRPV1 activation induced by the trigeminal agonist. This suggests that trigeminal activation can alter odorant responses even at the earliest stage of the olfactory sensory transduction.

**Significance Statement:** Most odorants reaching the olfactory epithelium can simultaneously activate olfactory and trigeminal systems. Although these two systems constitute two separate sensory modalities, trigeminal activation can alter odor perception. Here, we analyzed the trigeminal activity induced by different odorants proposing an objective quantification of their trigeminal potency independent from human perception. We show that trigeminal activation by odorants reduces the olfactory response in the olfactory epithelium and that such modulation correlates with the trigeminal potency of the trigeminal agonist. These results show that the trigeminal system impacts the olfactory response from its earliest stage.

## Introduction

Airborne chemicals are detected by olfactory sensory neurons (OSNs) in the olfactory epithelium (OE). The signal is peripherally transduced into action potentials, conveyed to the olfactory bulb (OB), and further processed and transmitted to cortical areas. Most studies of the olfactory system consider the OB as the first station of modulation of olfactory information (Schmidt and Strowbridge, 2014; Liu et al., 2015; Brunert and Rothermel, 2021). Few studies have explored modulation in the OE (Bouvet et al., 1988; Hegg et al., 2003; Daiber et al., 2013) although multiple systems could affect the olfactory signals in OSNs. The ethmoidal branch of the trigeminal nerve innervates both the OE and OB (Schaefer et al., 2002), and trigeminal-olfactory mutual modulation has been reported at the peripheral, central, and perceptual levels (Cain et al., 1980; Gudziol et al., 2001; Brand, 2006; Bensafi et al., 2007; Frasnelli et al., 2007; Lötsch et al., 2016; Tremblay and Frasnelli, 2018). fMRI studies showed cortical areas processing both nociceptive and olfactory stimuli (Bensafi et al., 2007; Lötsch et al., 2012; Pellegrino et al., 2017), while psychophysical studies demonstrated changes in trigeminal sensitivity influence the perception of odorants (Cain et al., 1980). The vast majority of odorants are also trigeminal agonists (Cometto-Muñiz and Cain, 1990; Cometto-Muñiz and Abraham, 2016). They typically activate the trigeminal system at medium to high concentrations, suggesting that when odorants enter the nasal cavity, both OSNs and trigeminal free-ending sensory fibers are activated (Doty et al., 1978; Cometto-Muñiz and Cain, 1990; Silver, 1992; Cometto-Muñiz and Abraham, 2016; Lötsch et al., 2016). When activated by odorants, different subsets of these trigeminal fibers will evoke specific sensations, described as pungent, tingling, stinging, burning, cooling, warming, painful, and irritating (Basbaum et al., 2009; Viana, 2011; Licon et al., 2018). Psychophysically, the trigeminal potency of odorants is described as the level of perceptual irritation they can evoke (Doty et al., 1978). Methods to determine the trigeminal potency of odors based on this definition are limited and provide only a subjective qualitative evaluation (Doty et al., 1978; Cometto-Muñiz et al., 2005). Currently, there is no quantitative parameter to complement such classification, highlighting the need analyze trigeminal neuronal responses to odorants.

Transient receptor potential cation channels (TRP channels), such as vanilloid 1 (TRPV1), ankyrin (TRPA1), and melastatin 8 (TRPM8), play key roles in the detection of odorants by the trigeminal system (Nilius and Owsianik, 2011; Nguyen et al., 2017). TRPV1 and TRPM8 are largely expressed on different subsets of trigeminal sensory fibers, except for a small population of TRPM8-expressing neurons that express TRPV1 as well (Hjerling-Leffler et al., 2007; Takashima et al., 2007; Huang et al., 2012; Nguyen et al., 2017). While TRPA1 is mostly co-expressed with TRPV1, a population of trigeminal sensory neurons express only TRPV1 (Bautista et al., 2005; Kobayashi et al., 2005; Nguyen et al., 2017; Yang et al., 2022). TRPA1 and TRPV1-expressing sensory fibers are also peptidergic and, when activated, release neuropeptides, such as calcitonin gene-related peptide (CGRP) and ATP into the surrounding epithelium (Holzer, 1998; Ding et al., 2000; Fabbretti et al., 2006; Shevel, 2014). The nasal mucosa, including the OE, is extensively innervated by peptidergic trigeminal fibers, where they can be detected alongside OSNs (Schaefer et al., 2002; Silver and Finger, 2009; Daiber et al., 2013). Previous work has shown a reduction of the OE response to odorants during the application of CGRP or ATP (Hegg et al., 2003; Daiber et al., 2013). We asked whether trigeminal activation modulates OSN activity in the OE. We show that activation of TRPA1 and TRPV1 channels by odorants reduces OSN responses. The stronger the trigeminal potency of an odorant, the greater the inhibition of the olfactory response. This suggests that trigeminal fibers can regulate the odorant response at its earliest stage, within the OE.

## Material and Methods

### Animals and ethical approval

C57BL6/J mice (purchased from The Jackson Laboratory, Bar Harbor, ME) were used as wild-type mice. TrpA1/V1-double KO mice, on a C57BL6 background (TRPA1/V1-KO), were a generous gift from Dr. Diana Bautista, University of California Berkeley (Gerhold and Bautista, 2008). In these mice, exon 23 (residues 901–951), which encodes the putative pore, and part of the sixth transmembrane domain of the TRPA1 receptor, is deleted (Bautista et al., 2006), as well as the fifth and all of the sixth putative transmembrane domains and the pore-loop domain of the TRPV1 receptor (Caterina et al., 2000). All animals were bred and housed in the animal facility of the Monell Chemical Senses Center in conventional polycarbonate caging with wood chip bedding (Aspen). Animals were kept at a 12-h light/dark cycle and *ad libitum* access to food and water.

All experimental procedures were performed in accordance with the National Institutes of Health (NIH) Guidelines for the Care and Use of animals and approved by the Monell Chemical Senses Center Animal Care and Use Committee. Every effort was made to minimize the number of animals used and their suffering.

### Primary trigeminal culture

Animals of both strains were euthanized by CO_2_, followed by cervical dislocation. The trigeminal ganglia from 3-4 mouse neonates (P3-P9) were surgically removed and transferred into Ca^2+^ and Mg^2+^ free Hanks’ Balanced Salt Solution (HBSS) including 1% penicillin/streptomycin (PS, 100 IU, and 100 μg/ml). Trigeminal ganglia were finely triturated, transferred into a 15 ml tube, and incubated in 5 ml 0.05 % trypsin Ca^2+^ and Mg^2+^ free HBSS-PS solution for 10 minutes at 37°C. 5 ml HBSS-PS solution was then added to stop active trypsin and centrifuged for 3 minutes at 300 x g. The supernatant was carefully discarded. After that, the TGNs were incubated in 5 ml 0.05 % collagenase A HBSS-PS solution for 20 minutes at room temperature, and 5 ml HBSS-PS solution was added and centrifuged for 3 minutes at 300 x g. The supernatant was discarded. 1 ml DMEM was added into the tube and triturated about 10-20 times at moderate force with a fire-polished pipette and seeded onto No#1 15 mm round coverslips coated by ConA at 37°C overnight.

### Ca^2+^ imaging

For our experiments, we used five different stimuli: 2-Phenylethanol (PEA), Pentyl Acetate (PA), Cinnamaldehyde (CNA), Allyl-isothiocyanate (AITC), Menthol (MNT), and Capsaicin (CAP). All were purchased from Sigma Aldrich. Cellular responses to these odorants were measured using a ratiometric Ca^2+^ imaging technique as previously described (Gomez et al., 2005). The cells were loaded with 5 μM acetoxymethyl-ester of Fura-2 (Fura-2/AM) and 80 μg /ml pluronic F127 (Molecular Probes, Eugene, Oregon) for at least 30 minutes at room temperature, settled in a recording chamber and superfused with Ringer’s solution or Ringer’s solution containing different chemical compounds via a valve controller (VC-8, Warner, USA) and perfusion pump (Perimax 12, SPETEC, Germany). Stimulation and washout duration was 20 - 30 s and 10 minutes at 3 ml/min perfusion rate, respectively, which depends on the chemical characteristics of the applied compound. There was a 10 s delay between solenoid valve activation and the arrival of stimulus compounds at the TGNs. Ca^2+^ imaging recordings were obtained using a Zeiss microscope equipped with a MicroMax RS camera (Roper Scientific Inc. Tuscon, AZ) and a Lambda 10-2 optical control system (Sutter Instrument Co. Novato, CA). Excitation from a monochromator was set at 340 nm and 380 nm with a 510 nm emission filter and the cellular fluorescence was imaged with a 10x objective (Zeiss). Images were digitized and analyzed using MetaFluor software (Molecular Devices, Sunnyvale, CA). Among the multiple types of cells in the dissociated tissue preparation, TGNs were recognized based on morphology and positive response to 30 mM KCl. To compare trigeminal potency across stimuli, we chose concentrations based on EC50 determined in previous literature (Bandell et al., 2004; Jordt et al., 2004; Bautista et al., 2007; Elokely et al., 2016; Lieder et al., 2020; Xu et al., 2020). For PA and PEA, which were not previously characterized, we used concentrations of similar magnitude to their olfactory EC50. The change in fluorescence ratio (F_340_/F_380_) was calculated for region-of-interests (ROIs) drawn manually around these cells. Response magnitudes were measured as the difference between the peak magnitudes (F_peak_) during the response window (90 s following presentation of stimulus) minus the mean baseline fluorescence (F_0_) and then divided by the mean baseline fluorescence ((F_peak_-F_0_)/F_0_). Increases of intracellular Ca^2+^ greater than 3% from the baseline level of fluorescence were considered as responses.

### Electro-olfactogram (EOG)

12 - 24 week old mice were euthanized by intraperitoneal injection of urethane (8mg/g of body weight, ethyl carbamate, Sigma Aldrich) followed by decapitation. We removed skin and lower jaw, and split the skull and nasal bone along the interfrontal and the internasal sutures. The olfactory endoturbinates were then exposed by removing the nasal septum.

We used the electro-olfactogram (EOG) set up and procedure similar to one previously described by Cygnar et al (Cygnar et al., 2010). The half head was mounted in an interface chamber with the sensory surface in constant contact with a stream of deodorized, humidified air, at a flow rate of 3 L/min. For each odorant (same as above) we prepared 5 M stock solution in DMSO (Sigma Aldrich), with the exception of MNT stock solution, which, due to its low solubility, was diluted to a 1 M stock. For dose-response experiments we used 10^-1^ to 10^-7^ serial dilutions of the stock solutions into water. Solutions were stored in glass vials with silicone stoppers and left to equilibrate with the air headspace for at least 30 min before the experiments. As a pure irritant stimulus we used a CO_2_/air mixture (50% v/v), which was prepared using a gas proportioner multitube flowmeter (Cole-Palmer, Vernon Hills, IL, USA), and stored in a sealed glass flask sealed during the experiment. For stimulation, odorants or CO_2_/air mixture were injected into the air stream with pressure pulses (100 ms, 10 psi) using a pneumatic picospritzer system (Parker Hannifin, Cleveland, OH, USA). Stimuli were presented at 1 minute intervals to allow the recovery of the epithelium.

Surface potentials from the endoturbinates 2 and 2b were recorded using two recording electrodes filled with 0.05% agarose melted in Ringer’s solution (mM): 140 NaCl, 5 KCl, 2 CaCl_2_, 1 MgCl_2_, 10 HEPES; (7.4 pH). Recording pipettes were pulled from borosilicate glass capillaries (o.d. 1.5 mm, i.d. 0.87 mm) to a tip aperture of 20–25 μm using a Flaming–Brown puller (Sutter Instruments, Novato, CA, USA). The recording electrodes and a ground electrode were connected to two Warner DP-301 amplifiers. The 1 kHz low-pass signal was digitized (CED Micro 1401 mkII digitizer) and processed by a PC. Signal acquisition software (Cambridge Electronic Design) was used to acquire the data at a sampling rate of 2 kHz.

### Data Analysis

Recordings were analyzed using Origin (8.5v, OriginLab Corp., Northampton, MA, USA). Recording baselines were determined as the potential values before each odor stimulus, and were subtracted from the trace. Net amplitudes of odorant-induced signals were grouped according to the mice genotypes and stimulus, averages for each group presented with the standard error of the mean (SEM). Statistical analysis was carried out using Jamovi (Version 1.6, the Jamovi project 2021, https://www.jamovi.org). Indicated significance levels were calculated using Welch’s unpaired t-test, One-Way ANOVA (Welch’s) with Tukey Post-Hoc Test, and Kruskal-Wallis non parametric One-Way ANOVA with Dwass-Steel-Critchlow-Fligner Post-Hoc tests as indicated. P < 0.05 (*), P < 0.01 (**), or P < 0.001 (***).

## Results

### Trigeminal responses to odorants

Trigeminal responses to odorants were assessed using Ca^2+^ imaging (Fig. 1A and B). We imaged 1386 TGNs from WT and 1683 TGNs from TRPA1/V1-KO mice responding to five odorants: 2-phenylethanol (PEA), pentyl acetate (PA), cinnamaldehyde (CNA), allyl-isothiocyanate (AITC) and menthol (MNT). All odorants tested except PEA are previously characterized TRPA1 or TRPM8 agonists (Peier et al., 2002; Bandell et al., 2004; Bautista et al., 2005; Richards et al., 2010). In addition, we used capsaicin (CAP) to evaluate the population of TRPV1-expressing neurons (Caterina et al., 1997; Silver et al., 2006). In WT mice, 302/1386 (21.78%) responded to any odorant, while in TRPA1/V1-KO only 115/1683 (6.83 %) TGNs responded to the chemosensory stimuli we tested (Fig. 1C). In WT, CAP activated the largest population of neurons (17.3 %), followed by PA, PEA, MNT, AITC and CNA.

**Figure 1:**
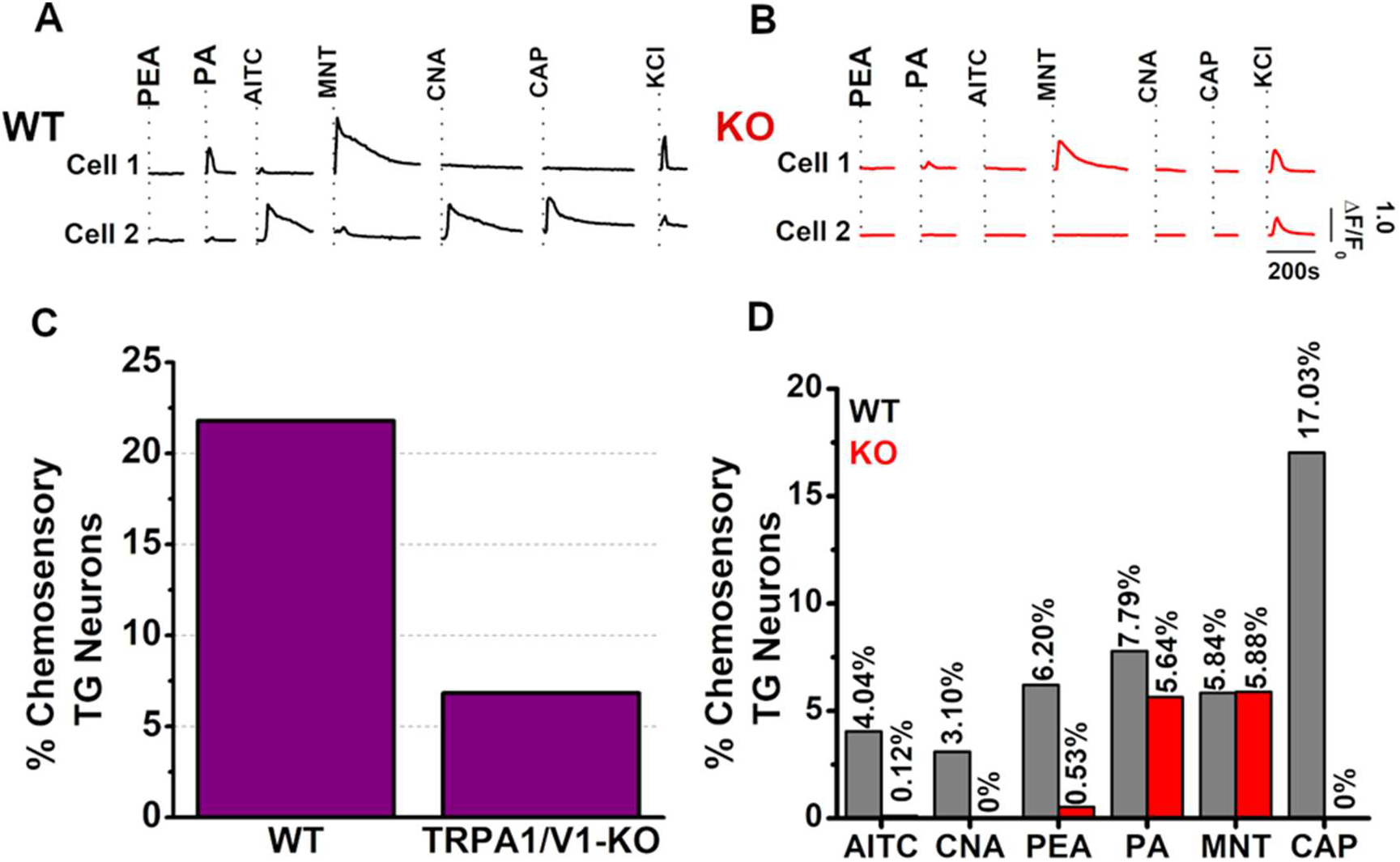
A, B) Example Ca^2+^ transients evoked by PEA (10 mM), PA (5 mM), AITC (50 μM), MNT (50 μM), CNA (400 μM), CAP (100 nM) and KCl in different cells in WT and TRPA1/V1-KO. C) Percentage of TGNs responding to odorants among all neurons obtained in the primary culture in WT (302/1386, 21.8%) and TRPA1/V1-KO (115/1683, 6.83%). D) Rate of TGNs responding to each stimulus in WT (grey) and TRPA1/V1-KO (red).

Overall, TGN primary cultures obtained from TRPA1/V1-KO mice show a lower percentage of neurons responding to chemosensory stimuli, with TGN responses to CAP, AITC, CNA and PEA being drastically reduced, while we could still record responses to MNT and PA (Fig. 1D). This suggests that TRPA1 and TRPV1 channels are necessary to evoke Ca^2+^ increases in response to AITC, CNA, CAP, and PEA. Furthermore, our results indicate that PA detection by the trigeminal system relies only partially on TRPA1 and TRPV1 channels, while MNT detection does not involve these two TRP channels.

For each odorant, we then obtained dose-response curves in the responsive subset of TGNs (Fig. 2). Maximal response amplitudes (V_max_) to AITC, CNA, and PA were significantly reduced in TRPA1/V1-KO (Welch’s unpaired t-test, AITC: df=19,p<0.001; CNA: df=52, p<0.001; PA: df=183, p=0.006), while no changes were observed for MNT and PEA (Welch’s unpaired t-test, PEA: df=66.6, p=0.06; MNT: df=134, p=0.313). In WT, MNT had the lowest EC_50_ (13.86 ± 2.70 μM, n=42), followed by AITC (61.63 ± 13.28 μM, n=20), CNA (0.264 ± 0.08 mM, n=53), PA (2.34 ± 0.55 mM, n=70) and PEA (8.53 ± 1.66 mM, n=35). In TRPA1/V1-KO, EC_50_ of MNT (10.14 ± 1.85 μM, n=94) and PA (11.67 ± 0.87 mM, n=128) showed no significant changes (Welch’s unpaired t-test, MNT: df=80, p=0.26; PA: df=192, p=0.35). TGNs of TRPA1-V1-KO responding to PEA showed decreased sensitivity to the odorant (77.77 ± 19.3 M, n=46, Welch’s unpaired t-test, p<0.001, F=45), while AITC and CNA did not evoke any response in TRPA1/V1-KO. Overall, we observed changes in either EC50 or V_max_, for all odorants except MNT, suggesting no involvement of TRPA1 or TRPV1 in its detection by the trigeminal system.

**Figure 2:**
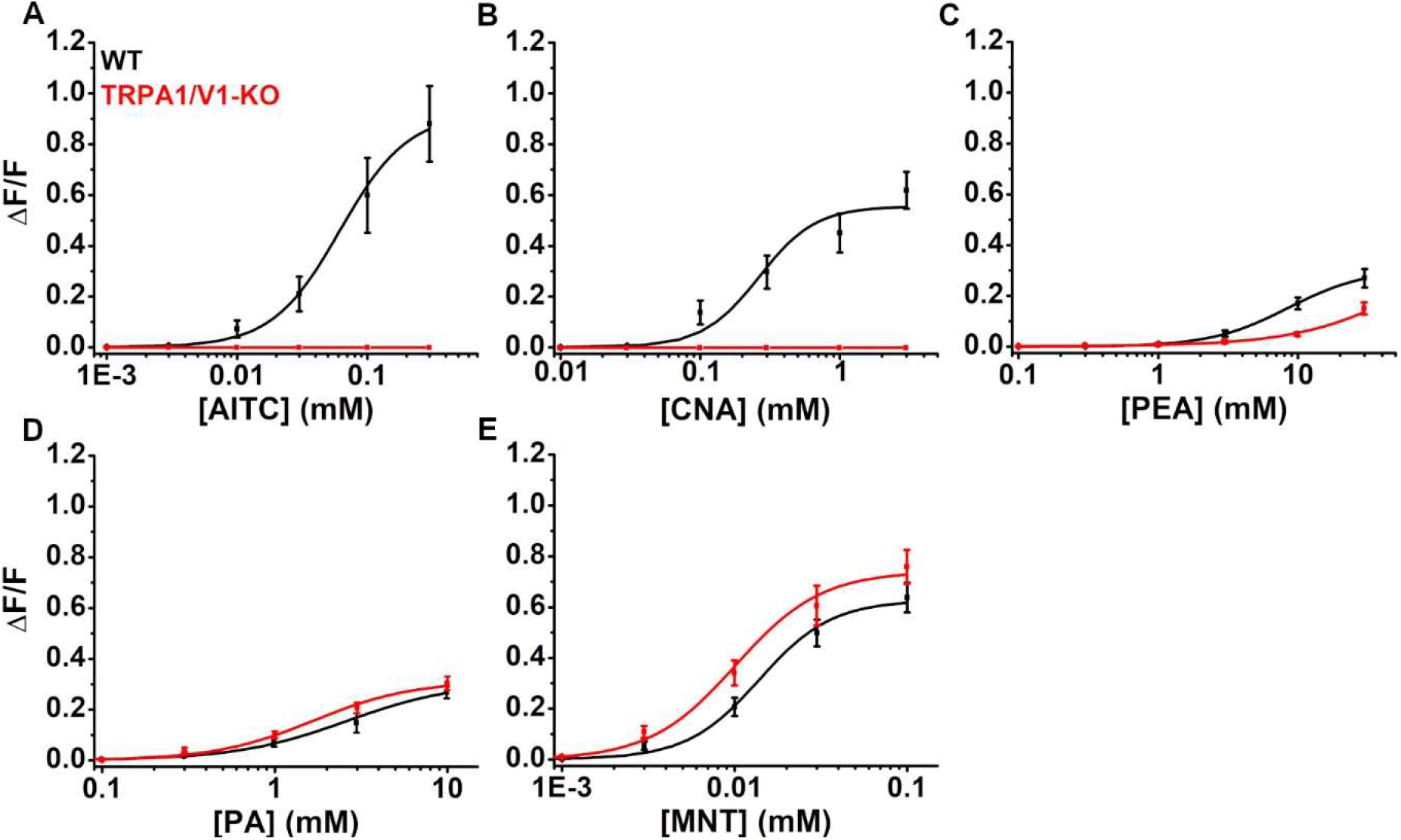
dose-response curves from TGNs in WT (black) and TRPA1/V1-KO (red) for the odorants A) AITC; B) CNA; C) PEA; D) PA; E) MNT.

### Physiological classification of trigeminal activation by odorants

To quantified the activity evoked by odorants in single cells expressing a given receptor we used an “activity index”, which was defined as (−log(EC_50_ (mM)) × max ΔF/F) (del Mármol et al., 2021). We used a similar approach to quantify the overall activity induced by each agonist across the population of TGNs, we multiplied the activity index of each odorant in each mouse strain by the percentage of TGNs they activated (Fig. 1D), generating a “weighted” activity index (WAI, Fig. 3B). AITC and CNA activity index in TRPA1/V1-KOs is 0, due to the lack of responses to these odorants in the absence of TRPA1 and TRPV1 channels. Finally, to summarize the quantitative trigeminal properties of an odorant, we subtracted WAI_KO_ from WAI_WT_ to obtain a single score (TRPA1/V1 score) for each odorant (Fig. 3C). Positive scores correspond to predominantly TRPA1 and TRPV1 agonists, like AITC, CNA and PEA. PA and MNT trigeminal scores have negative values, reflecting an increase of trigeminal activity evoked by these two odorants in TRPA1/V1-KO, which is not mediated by these two TRP channels.

**Figure 3:**
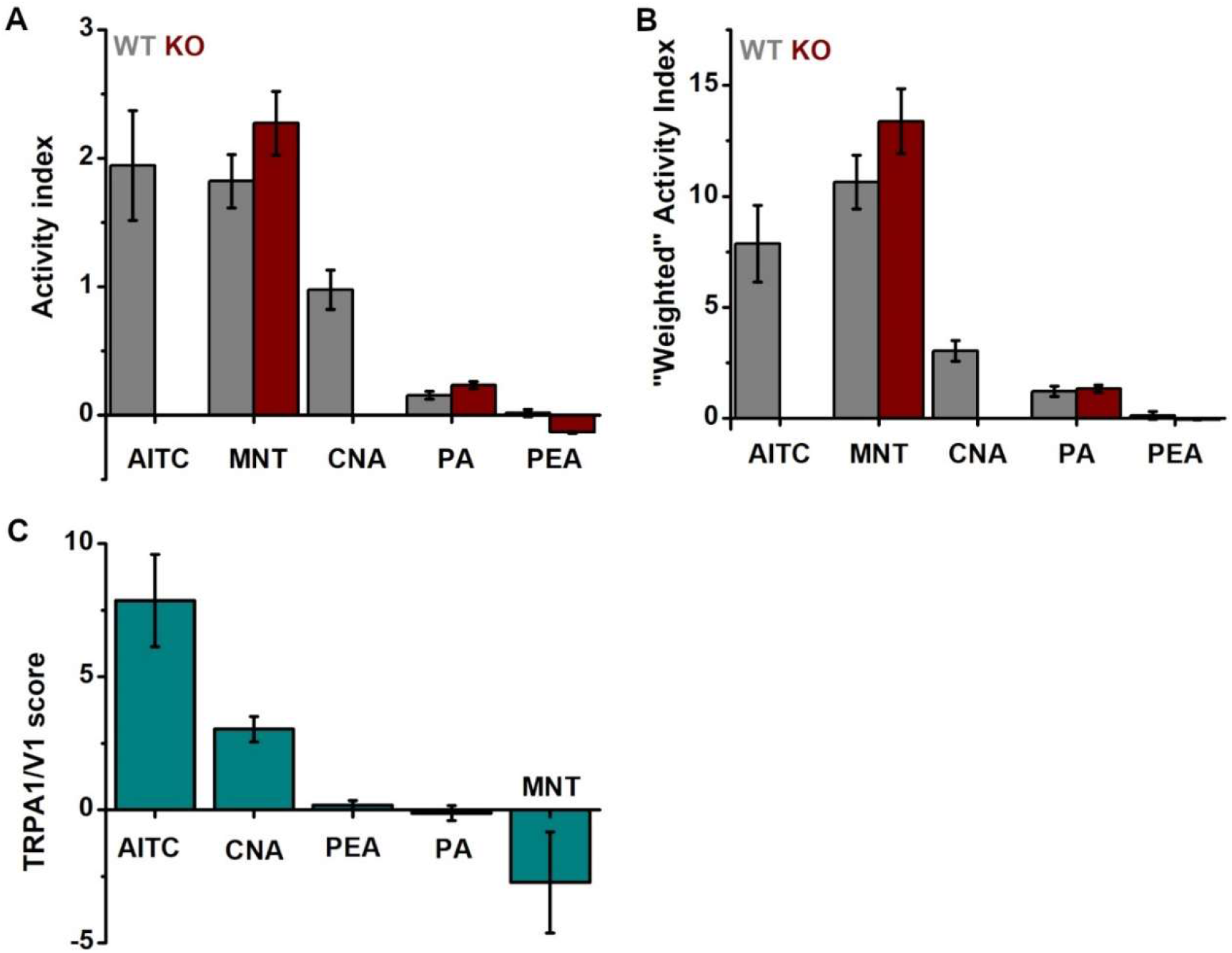
A) Activity index of all odorants in WT (grey), and TRPA1/V1-KO (red). B) Weighted activity index of all odorants in WT (grey), and TRPA1/V1-KO (red). C) TRPA1/V1 score, positive values are associated with odorants that are predominantly TRPA1/V1 agonists (AITC, CNA and PEA), while negative scores are associated with odorants which activate predominantly other TRP receptors (PA and MNT).

### Odorant responses in TRPA1/V1_KO mice

Using the EOG technique, we assessed if the lack of expression of TRPA1 and TRPV1 receptors in the OE could affect the response to odorants. We determined dose-response relations in the OE for all the odorants previously tested on TGNs (except CAP). Peak response amplitudes for each concentration were fitted with a Hill equation to obtain a dose-response curve. CNA, MNT and PEA showed no significant differences among dose-response curves obtained in WT and TRPA1/V1-KO (Fig. 4). The absence of TRPA1 and TRPV1 channels in the KO mice resulted in a leftward shift of the dose response curve of AITC and PA, both characterized by a significant reduction of Ks (Fig. 4L; AITC: F(1,7.27)=4.73, p=0.047; PA: F(1,7.28), p=0.19; Welch’s One-Way ANOVA, Tukey Post-Hoc Test) and therefore sensitization to odorants. Maximal response amplitudes (V_max_) were unchanged in WT and TRPA1/V1-KO across all odorants (Fig. 4K).

**Figure 4:**
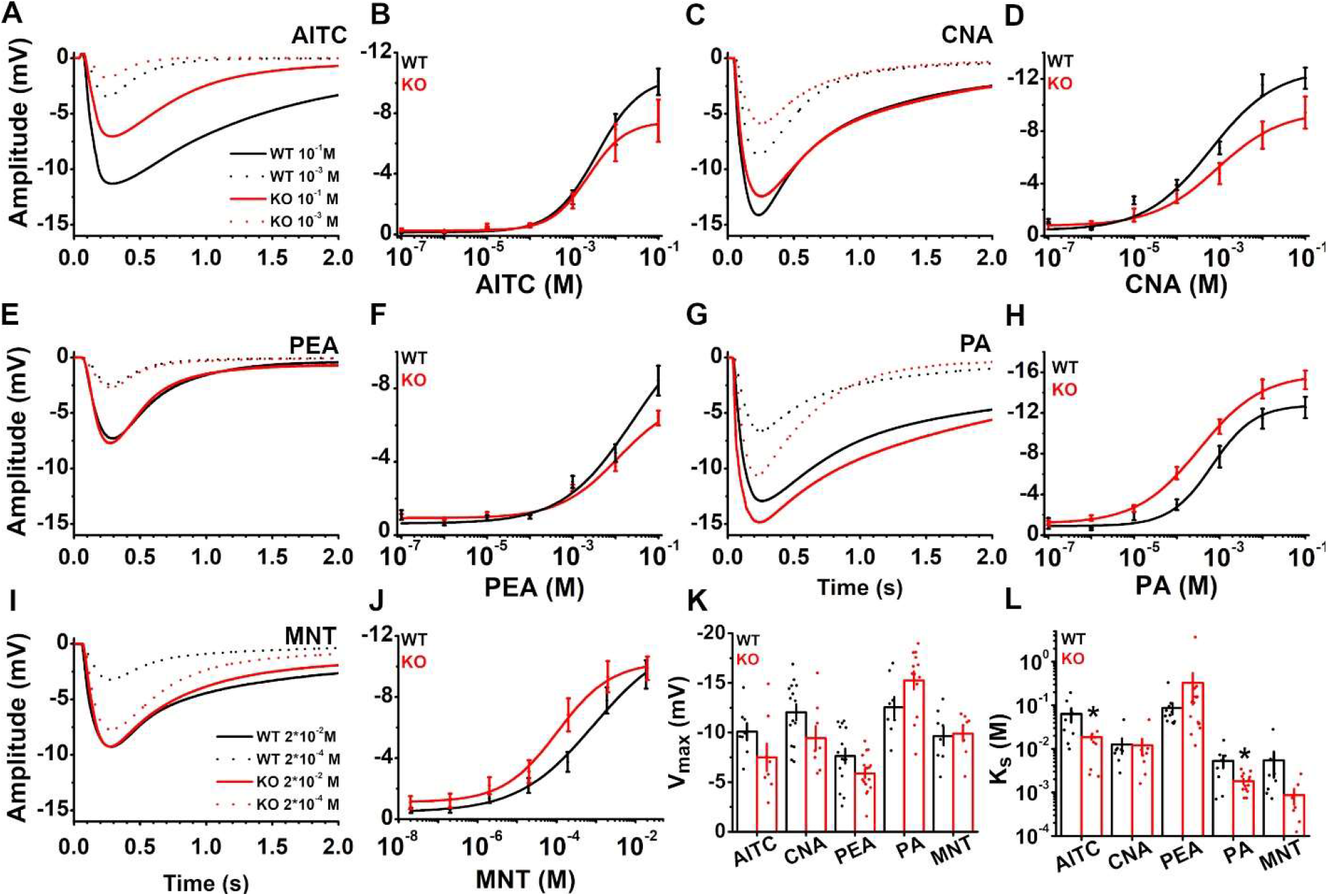
A, C, E, F, I) Examples of EOG responses in WT (black) and TRPA1/V1-KO (red) to two different concentrations (10^-2^ and 10^-4^) of odorants. B, D, F, H, J). Dose-response curves of all odorant in WT (black) and TRPA1/V1-KO (red). K, L) Maximal response amplitude and K_s_ obtained for all odorants in WT (red) and TRPA1/V1-KO (red). Significance was calculated using a Welch’s one-way ANOVA with Tukey Post-Hoc Test. WT vs TRPA1/V1-KO. P < 0.05 (*).

### Repeated exposure of the OE to irritants reduces the EOG response to odorants

To establish if trigeminal activation by odorants can modulate the olfactory response, we performed EOG recordings, in which we alternated brief stimulations of the OE with PEA (0.1 M), the odorant with the lowest trigeminal potency, followed by exposure of the OE to a trigeminal agonist to activate the trigeminal sensory fibers. We applied three pulses of PEA (100 ms) to the OE alternated with pulses of a given trigeminal agonist (100 ms), followed by three more PEA pulses (Fig. 5A). In between each stimulus we allowed 1 min for the OSNs to recover from the previous stimulation and to avoid olfactory adaptation. To determine how the response to PEA would change during the recording session, in the absence of any trigeminal stimulus, we alternated PEA and a pulse of non-odorized air. EOG responses to PEA were normalized by dividing each EOG peak amplitudes (V) by the amplitude of the first PEA response (V_0_). The means of the normalized EOG responses were then compared (Fig.5D and E). In the control, we observed a decrease of the PEA response amplitude during the experiment reaching approximately 22 % in the last PEA stimulus (Fig. 5B, D). This decline was observed in both WT and TRPA1/V1-KO. We then repeated the same experiment using CO_2_ (50 % v/v, 100 ms pulse), a potent TRPA1 agonist (Fig. 5C, E). In WT, CO_2_ induced a reduction of the PEA response of approximately 53 % (Fig. 5E, black). Such stark reduction of the EOG response was not observed in TRPA1/V1-KO mice (Fig. 5E, red), in which the responses to PEA were no different from the control.

**Figure 5:**
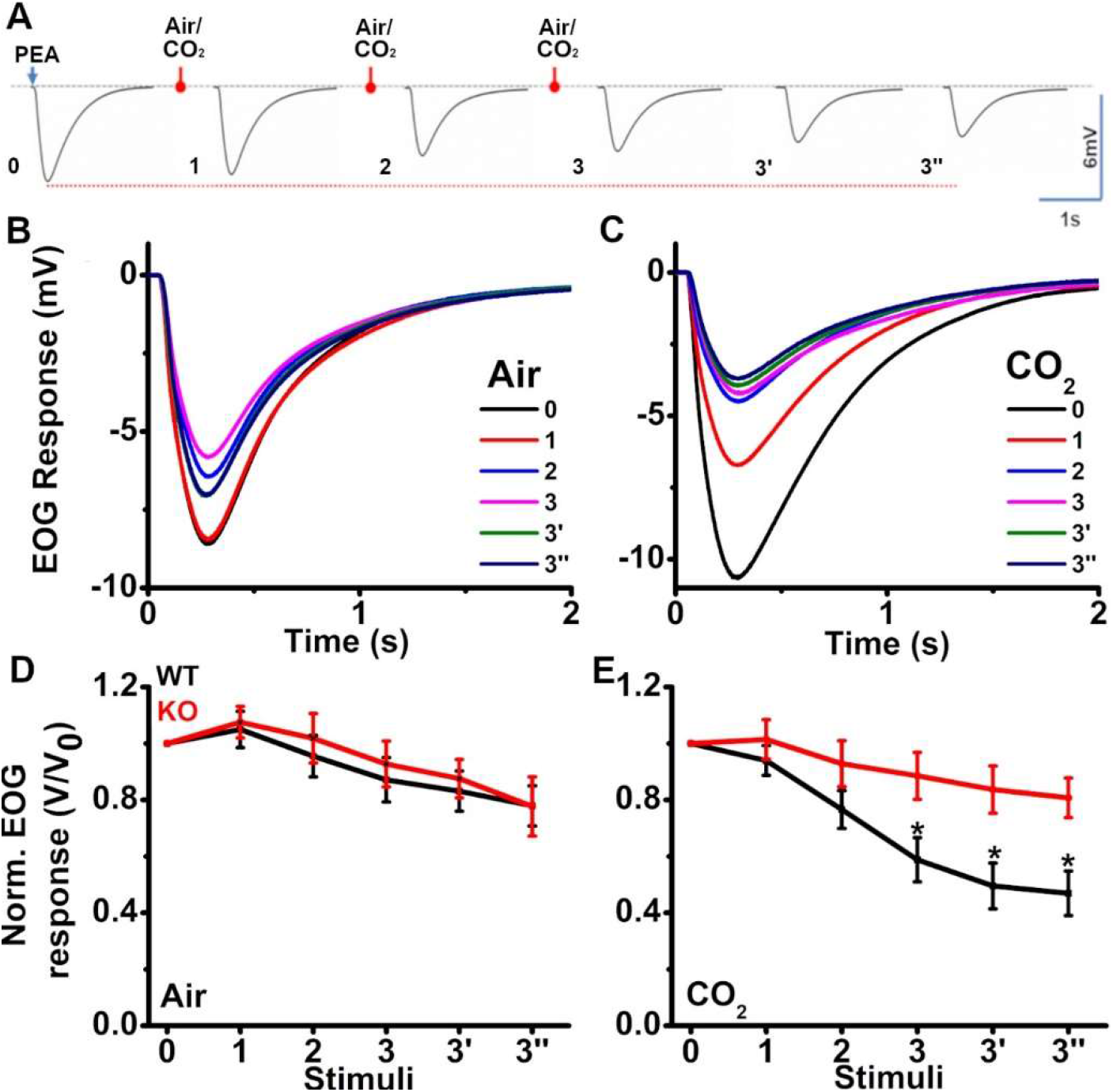
A) Sequence representing the experimental protocol. B and C) Example of EOG responses to PEA in WT when OE was exposed to air or CO_2_. D and E) Mean EOG responses to PEA normalized to the amplitude of the first PEA response (V_0_) in WT (black) and TRPA1/V1-KO (red) after exposure to air (D, WT n=19; KO n=7) or to CO_2_ (E, WT n=14; KO=12). Significance was calculated using Kruskal-Wallis non parametric One-Way ANOVA with Dwass-Steel-Critchlow-Fligner Post-Hoc Test. WT vs KO. P < 0.05 (*).

We then addressed if odorants which are trigeminal agonists could elicit the same modulation of the OE activity as CO_2_. AITC (0.1 M), which has the highest TRPA1/V1 score elicited a similar effect as CO_2_, inducing a progressive reduction of the PEA response in WT, which was abolished in TRPA1/V1-KO mice (Fig. 6A, Mean V_3”_/V_0_ WT: 0.36 ± 0.087, n=11; KO: 0.67 ± 0.076, n=10, p=0.024, Kruskal-Wallis non parametric One-Way ANOVA, with Dwass-Steel-Critchlow-Fligner Post-Hoc Test).

**Figure 6:**
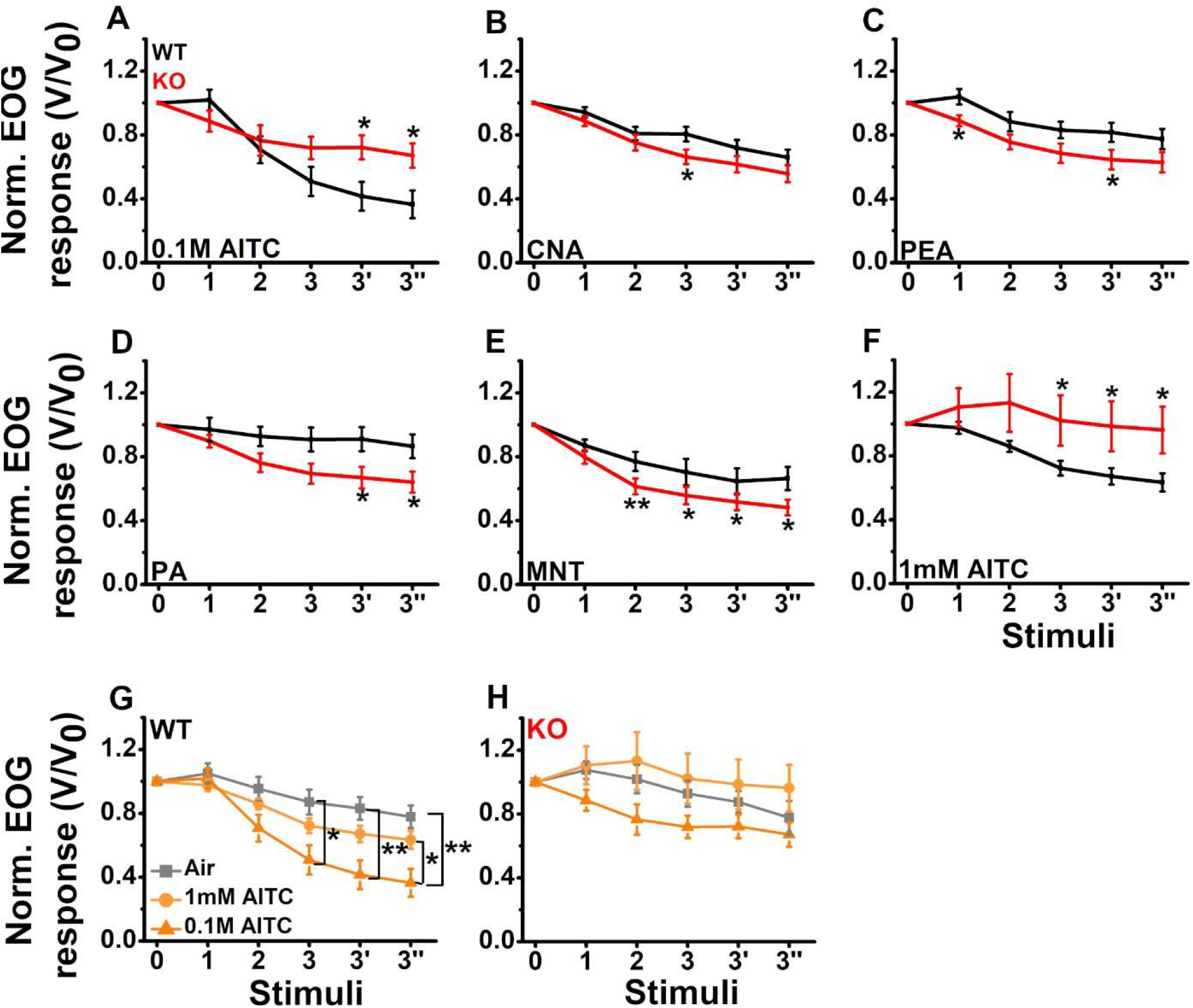
A-F) Mean EOG responses to PEA normalized to the amplitude of the first PEA response (V_0_) after exposure of the OE to 0.1M AITC (A), CNA (B), PEA(C), PA (D), MNT (E) and 1mM AITC (F). Comparison of mean normalized EOG responses to PEA in WT (G) and TRPA1/V1-KO (H) after the exposure of the OE to 0.1M AITC (triangle, dark orange lines), 1mM AITC (circles, light orange lines) and Air (squares, grey lines). Significance was calculated using Kruskal-Wallis non parametric One-Way ANOVA with Dwass-Steel-Critchlow-Fligner Post-Hoc Test. WT vs KO P < 0.05 (*), P < 0.01 (**).

CNA (0.1 M) and PEA (0.1 M) did not induce a difference among EOG responses in WT and TRPA1/V1-KO, which declined to the same rate by the end of the experimental protocol (Fig. 6B, C). In WT, we observed a small and temporary enhancement of the olfactory response after three CNA stimuli (WT: V_3_/V_0_: 0.81 ± 0.04, n=23; KO: 0.75 ± 0.05, n=22, p = 0.017). Similarly in WT, we observed the same enhancement of the relative response amplitude (V/V_0_) of the stimuli 1 and 3’, when PEA was used as the trigeminal agonist but the EOG responses in response to the olfactory stimuli thereafter (2 and 3’’) were not significantly different from TRPA1/V1-KO (Mean V_1_/V_0_ WT: 1.04 ± 0.0481, n=20; KO: 0.89 ± 0.03, n=13; p=0.011. Mean V_3’_/V_0_ WT: 0.81 ± 0.06, n=20; KO: 0.64 ± 0.06, n=13, p = 0.031). The stimulation of the OE by PA (0.1 M) and MNT (0.02 M) induced a more robust and sustained decay of the response to PEA in TRPA1/V1-KO (Fig. 6 D, E), which persisted until the end of the recordings (PA Mean V_3”_/V_0_ WT: 0.87 ± 0.07, n=18; KO: 0.64 ± 0.07, n=13, p = 0.038; MNT Mean V_3”_/V_0_ WT: 0.66 ± 0.07, n=16; KO: 0.48 ± 0.05, n=13, p = 0.02).

We then repeated the previous experiment exposing the OE to a lower concentration of AITC (1 mM) to test if the trigeminal modulation of the olfactory response is concentration dependent. Exposing the OE to 1 mM AITC still induced a significant reduction of the odor response in WT in comparison to TRPA1/V1-KO (Fig. 6F, Mean V_3”_/V_0_ WT: 0.63 ± 0.06, n=23; KO: 0.96 ± 0.15, n=11, p < 0.01), but significantly smaller in comparison to the one elicited by 0.1 M AITC in the same strain (p = 0.034), and not significantly different to the control with air (Fig. 6G). In TRPA1/V1-KO mice, neither concentration of AITC altered the PEA response relative to the control (Fig. 6F).

### TRPA1/V1-score correlates with the level of trigeminal modulation of OE response to odor in WT

We next determined if the ability of an odorant to activate TRPA1 and TRPV1-expressing trigeminal fibers correlates with the reduction of the olfactory response induced by the same odorant in the OE (Fig. 7). For both strains we plotted the reduction of the PEA response (%) induced by the odorant against its TRPA1/V1 score. MNT was excluded from this analysis as it does not activate TRPA1 or TRPV1 channels. TRPA1/V1 score of 1 mM AITC (0) was calculated based on the AITC dose response curve obtained in TGNs. All data points were fitted with a linear function (Fig. 7A, WT: intercept = 24.04 ± 5.75; slope = 4.48 ± 1.83; Fig. 7B, KO: intercept = 36.50 ± 6.26; slope = 0.25 ± 1.72). In WT, the reduction of the olfactory response correlates with the TRPA1/V1 score of the odorant (R = 0.817, p=0.091), while changes in odor responses in TRPA1/V1-KO are not linked to the trigeminal properties of the odor (R = 0.0838, p=0.89).

**Figure 7:**
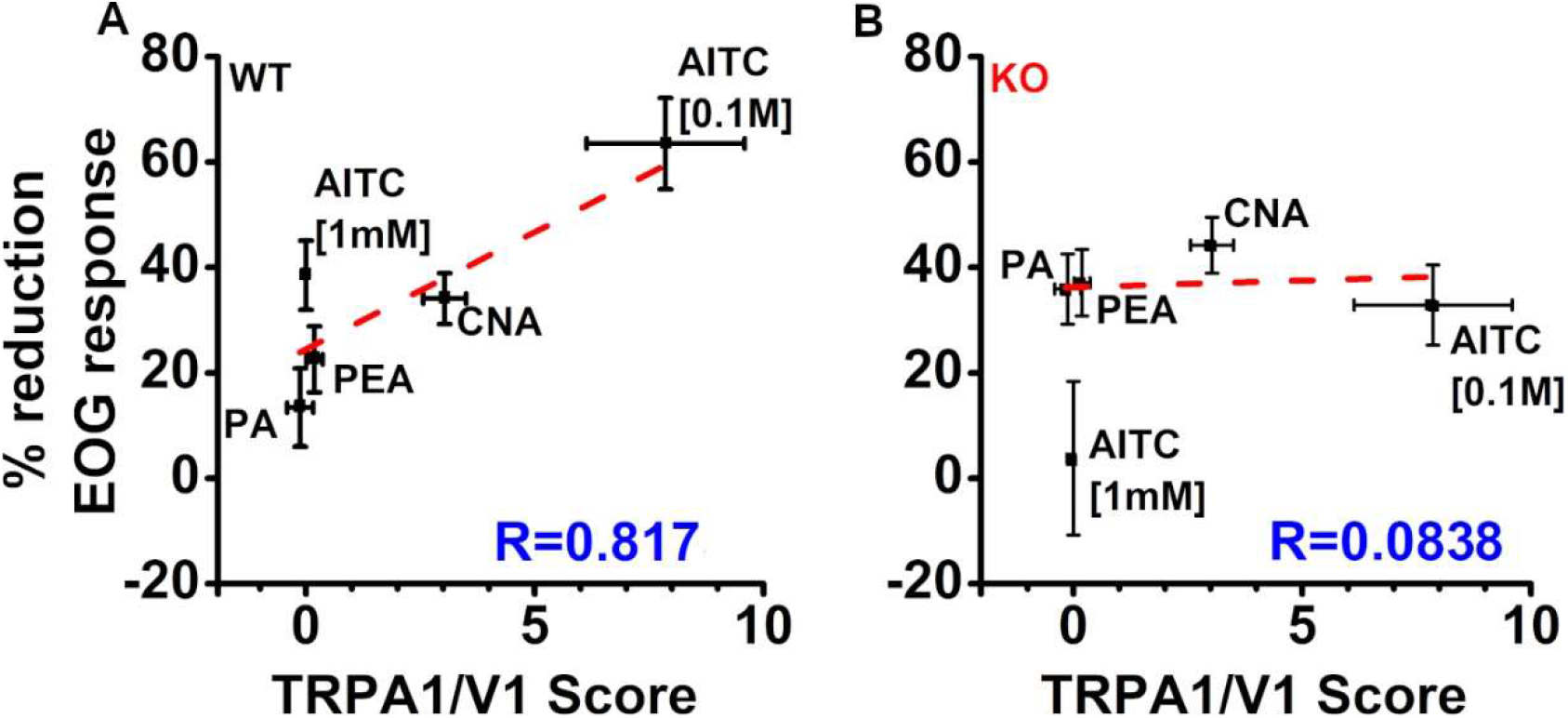
Correlation TRPA1/V1 score and reduction EOG response (%) in WT (A) and TRPA1/V1-KO (B)

## Discussion

In this work we quantified the trigeminal activity induced by odorants and how it modulates the olfactory response generated in the OE. Until now, trigeminal potency has been described using only psychophysical approaches (Doty, 1975; Doty et al., 1978; Cometto-Muñiz and Cain, 1990; Frasnelli and Hummel, 2007; Cometto-Muñiz and Abraham, 2016). With such methods it is hard to separate the two sensory modalities evoked by the odorant, and they provide only a subjective evaluation of the perception evoked. Assessments of trigeminal potency of odors by patients with olfactory loss eliminates the olfactory interference from the measurements, but acquired anosmia is associated with an alteration of trigeminal perception as well (Gudziol et al., 2001). While more objective methods to measure trigeminal responses to odorants from patients like the recording of the negative mucosal potential and functional magnetic resonance are less suitable for screenings on a large scale (Kratskin et al., 2000; Bensafi et al., 2012; Pellegrino et al., 2017). Based on the responses to odorants obtained in TGNs we developed the TRPA1/V1-score, a physiological classification of trigeminal potency of odorants. For each odorant, this score incorporates their activity index, the size of the trigeminal population activated by it and if it activates TRPA1 and/or TRPV1 channels. Although this score does not provide a further distinction among different chemosensory TRP channels, it is the first quantitative measure of trigeminal potency without any olfactory interference and independently from human perception. Previously, a few works have suggested the possibility of trigeminal/olfactory interaction at the periphery. Tracing of the trigeminal innervation of the nasal cavity showed previously that peptidergic sensory fibers from the ethmoidal branch of the trigeminal nerve innervate the OE and OB (Finger and Böttger, 1993; Schaefer et al., 2002).

Previous work from Hegg et al. and Daiber et al. showed that ATP and the neuropeptide CGRP can both modulate OSN responses to odorants (Hegg et al., 2003; Daiber et al., 2013). Both compounds are released upon stimulation by trigeminal sensory fibers, which express TRPA1 and TRPV1 channels. Our work directly builds on Daiber’s and Hegg’s findings (Hegg et al., 2003; Daiber et al., 2013), addressing if the exposure of the OE to odorants with different trigeminal potencies could modulate the olfactory response, and if different levels of trigeminal activation would affect the olfactory response differently. Our results suggest that TRPA1/V1-agonists induce a graded modulation of the olfactory response to PEA, which correlates with the level of trigeminal activation they induce. This correlation is no longer present in the absence of TRPA1 and TRPV1 expression, suggesting that the TRPA1/V1-score is a valid indicator of trigeminal potency.

This modulatory mechanism likely originates from trigeminal sensory fibers rather than other cell types in the OE. Single-cell RNA-seq obtained from the OE shows a lack of expression of TRPA1 in non-neuronal cells (Tsukahara et al., 2021). Low levels of expression of TRPV1 have been detected in Trpm5^+^/Chat^+^ microvillar cells, but the involvement of these cell types in the modulation seems unlikely, since AITC and CO_2_ are both TRPA1, and not TRPV1 agonists.

A subpopulation of trigeminal TRPA1/V1-expressing fibers are peptidergic free nerve endings, which, when stimulated can release neuropeptides such as CGRP or neuromodulators like ATP. The activation of this population of sensory neurons by odorants might induce the release of different amounts of ATP and CGRP, as measured by the score TRPA1/V1 score. Previous studies have shown that ATP and CGRP can reduce the olfactory response in the OE (Hegg et al., 2003; Daiber et al., 2013), possibly driving Ca^2+^ in the dendritic and soma compartments. The increase of intracellular Ca^2+^ could then activate Ca^2+^-activated K^+^ currents (Kawai, 2002) and, consequently, decrease OSN responses evoked by the following stimulus. OSNs express purinergic receptors P2X4 and P2Y2 (Xu et al., 2016; Tsukahara et al., 2021). The release of ATP into the extracellular space could open P2X4 expressed on the membrane of OSNs and drive an intracellular Ca^2+^ increase (Stokes et al., 2017). Activation of P2Y2 receptors in OSNs would initiate the PLC-mediated Ca^2+^ signaling cascade, which leads to the release of Ca^2+^ from intracellular stores. Purinergically-induced intracellular Ca^2+^ increase in the OSNs might therefore contain two phases, an early one, driven by P2X4 and a delayed one, mediated by P2Y2. ATP in the intracellular space is quickly degraded, therefore combining both P2X and P2Y receptors might be crucial to provide a more sustained Ca^2+^ increase able to affect the odor response. The neuropeptide CGRP, which is also released by trigeminal peptidergic fibers, was also shown to modulate OSN responses to odorants (Daiber et al., 2013). The activation of the CGRP receptor leads to the activation of adenylate cyclase followed by an increase of cAMP (Russell, 2011), and consequentially to the rise of intracellular levels of Ca^2+^. In the OSNs CGRP has been shown to induce increases in cAMP (Daiber et al., 2013), which could contribute to drive intracellular Ca^2+^ increase and affect their response to odorants.

Taken together our study shows that odorants can simultaneously activate both the olfactory and trigeminal system in the OE, and that TRPA1/V1-extressing trigeminal fibers can modulate the OSN response to odors. Such modulation is a graded reduction of the olfactory signal which correlates with the odorant’s TRPA1/V1-score. The TRPA1/V1-score we developed can predict the impact that previous exposure to TRPA1/V1-agonists can have on OSN activity, but further studies, and more odorants will need to be tested to determine to what extent trigeminally active odorants can affect the OSNs responses. The mechanism we describe supports and complements the previous findings of a peripheral modulation of the olfactory signal by the trigeminal system and underscores the necessity of taking into account the trigeminal potency of an odorant when analyzing olfactory sensory processing. Furthermore, the role of the trigeminal activation might be particularly relevant when considering odor mixtures coding, with more than one component able to simultaneously activate the trigeminal system.

## Acknowledgments

This work was supported by NIH NIDCD R21DC018358 to FG, R01DC016598, R03DC012413, R01DE028979 to MT, and 1R01DC016647 to JR.

We thank Kevin Bolding and Joel Mainland for discussing the manuscript, and Minliang Zhou for the help with the animal colonies.

Marco Tizzano’s current affiliation is the University of Pennsylvania, Department of basic and translational sciences, 19104 Philadelphia PA, USA.

## Notes

Conflict of interest: The authors declare no competing financial interests.

### Competing Interest Statement

The authors have declared no competing interest.

